# Proteome Profiling of Human Tear Fluid Following Acute Exercise

**DOI:** 10.64898/2026.08.12.744559

**Authors:** Mingjie Sun, Huilin Yao, Maodi Liang, Qintong Fei, Jing Cao, Tiantian Liang, Qinghua Cui

## Abstract

Acute exercise induces systemic molecular perturbations that are not fully reflected by individual biomarkers. Tear fluid is amenable to non-invasive and repeated collection and contains a diverse array of bioactive molecules, thereby constituting a practical biological specimen for evaluating exercise-related physiological responses. However, the immediate proteome-wide alterations in tear fluid following acute exercise have not been characterized. In this study, we performed quantitative proteomic profiling of paired tear samples obtained from healthy female participants before and immediately after a single exercise session, using data-independent acquisition liquid chromatography–tandem mass spectrometry (DIA-LC– MS/MS). Among the 3,173 identified proteins, 744were significantly altered post-exercise, of which 484 exhibited up-regulation and 260 exhibited down-regulation. Functional enrichment analysis demonstrated that the up-regulated proteins were predominantly associated with translation and ribosome biogenesis, whereas down-regulated proteins were primarily involved in glycan metabolism, lysosomal processing, and extracellular matrix organization. Collectively, these observations indicate that acute exercise elicits a rapid and coordinated reconfiguration of the tear proteome. To our knowledge, this investigation provides the first proteomic evidence of the immediate tear response to physical activity, offering a molecular basis for understanding exercise-mediated modulation of tear composition and ocular surface homeostasis.

## 1. Introduction

Regular physical exercise confers substantial benefits for cardiometabolic function, physiological adaptability, and overall health^1–5^. Such benefits arise largely from cumulative adaptations to repeated bouts of exercise, each of which initiates rapid molecular responses that contribute to longer-term physiological adaptation. Acute exercise induces coordinated alterations in energy metabolism, circulating mediators, oxidative stress, inflammatory responses, and immune-related processes^6–8^. The complexity and interdependence of these responses limit the capacity of individual biomarkers to comprehensively represent exercise-induced physiological regulation. Molecular profiling of body fluids therefore provides a systematic approach for characterizing the early biological response to acute exercise.

Blood, saliva, and sweat have been extensively used to characterize exercise-induced alterations in metabolism, inflammation, oxidative stress, and immune regulation^9,10^; by comparison, the application of tear fluid in exercise research remains unavailable. Its collection is minimally invasive, requires only a small sample volume, and permits repeated sampling, providing practical advantages for monitoring molecular responses to physiological interventions. The molecular composition of this biofluid encompasses diverse bioactive constituents, including lacrimal gland-derived proteins, mucins, antimicrobial peptides, immunoglobulins, complement components, and extracellular vesicle-associated proteins^11,12^. Direct contact with the ocular surface enables its molecular profile to reflect changes within the local microenvironment. Furthermore, compositional alterations have been documented in association with ocular surface disorders, diabetes, autoimmune diseases, and neurodegenerative conditions^13–16^, supporting the capacity of tear fluid to reflect molecular changes related to both local ocular and systemic physiological or pathological states.

Despite this potential, existing studies of exercise and tear fluid have primarily evaluated tear secretion, tear-film stability, osmolarity, and selected inflammatory or oxidative stress markers^17^. Consequently, the extent to which acute exercise modifies the broader molecular composition of human tears remains unclear. ^11,12,18^. A proteome-wide analysis could therefore extend previous targeted observations by identifying coordinated changes in protein abundance and the biological processes underlying the immediate tear response to exercise.

Quantitative proteomics enables the simultaneous characterization of protein composition, abundance, and functional properties across biological conditions. Tear proteomics has been increasingly applied to ocular and systemic disease research, demonstrating its capacity to characterize molecular alterations within the ocular surface microenvironment^16,19,20^. However, the limited volume of tear samples and their broad protein abundance range impose substantial analytical demands. Liquid chromatography-tandem mass spectrometry (LC-MS/MS) operated in data-independent acquisition (DIA) mode provides extensive proteome coverage and reproducible quantification in small-volume samples, offering a suitable approach for the systematic investigation of exercise-associated changes in tear proteins^21,22^.

Against this background, the present study applied quantitative proteomic profiling to paired tear samples collected from healthy participants before and immediately after acute exercise. The study was designed to determine whether acute exercise induces detectable changes in the human tear proteome and to characterize the principal biological processes represented by these alterations. Establishing the immediate exercise-responsive tear proteomic profile provides a molecular reference for understanding exercise-associated changes in tear composition and ocular surface homeostasis under healthy conditions.

## 2. Materials and methods

### 2.1. Study design

Five healthy female participants, aged 23-27 years, with no history of exercise addiction or long-term professional training, were enrolled in this study. To minimize metabolic state variations caused by dietary differences, all participants were required to strictly adhere to a standardized diet for three days prior to the exercise intervention. This protocol prohibited the consumption of alcohol, coffee, strong tea, caffeine-containing beverages, as well as high-sugar, high-fat, and spicy stimulant foods. A moderate-intensity aerobic exercise protocol (treadmill running) was uniformly administered. Exercise intensity was precisely controlled using individual maximum heart rate (HRmax), calculated according to the Tanaka formula: HRmax = 208 – (0.7 × age)^23^. The target intensity range was set at 60%–80% of HRmax. The exercise session began with a 5-minute warm-up on the treadmill at 4 km/h, followed by 30 minutes of continuous running while maintaining the target heart rate zone. Heart rate was monitored in real-time throughout the session using a Polar H10 chest strap to ensure exercise intensity remained within the prescribed range. Tear fluid samples were collected immediately before and after the exercise session. The study protocol was approved by the Ethics Committee of the Wuhan Sports University (Approval No. 2025089), and written informed consent was obtained from all participants.

### 2.2. Collection and processing of tear fluid samples

Tear fluid samples were collected bilaterally using Schirmer test strips without local anesthesia, and sterile strips (5 mm wide) were placed at the outer one-third of the lower conjunctival sac until fully saturated. Tear-saturated strips were immediately transferred to a pre-cooled container and then stored at −80°C for subsequent analysis. For processing, samples were thawed at 4°C. Each test strip was then transferred to a microcentrifuge tube, and 400 μL of ice-cold methanol-water solution (4:1, v/v) was added. The mixture was subjected to ultrasonication in an ice bath for 30 minutes, followed by incubation at −20°C for 30 minutes. After centrifugation at 16,000 × g for 20 minutes at 4°C, the supernatant was collected for subsequent mass spectrometry analysis.

### 2.3. Proteomic analysis by data-independent acquisition mass spectrometry

Quantitative proteomic analysis was performed using DIA. Proteins extracted from samples were digested with trypsin. The resulting peptides were separated using a Vanquish Neo ultra-high-performance liquid chromatography (UHPLC) system (Thermo Scientific, Waltham, MA, USA) and analyzed by LC-MS/MS operated in DIA mode. During DIA acquisition, the full mass scan range was divided into consecutive windows, and all precursor ions within each window were cyclically fragmented to unbiasedly acquire fragment ion information for all detectable peptides. All raw MS data were processed, searched, and quantified using the DIA-NN software. The search was performed against the UniProtKB Homo sapiens (Human) protein database (version 20250414, downloaded from https://www.uniprot.org/taxonomy/9606). Protein quantification was based on the integrated peak areas of high-confidence peptide fragment ions.

### 2.4. Statistical analysis

All statistical analyses were performed using R (version 4.5.1). MS intensity values were retained for proteins quantified in at least 50% of samples; missing values were imputed using the K-nearest neighbor (KNN) method. Data were median-normalized and log₂-transformed. Differentially abundant proteins were defined as those with |log₂FC| > log₂(1.5) and P < 0.05.

GO and KEGG enrichment analyses were conducted using clusterProfiler and org.Hs.eg.db. GSEA was performed with MSigDB Hallmark gene sets using all quantified proteins ranked by post- versus pre-exercise log₂FC; enrichment was evaluated using the normalized enrichment score (NES), nominal P-value, and false discovery rate (FDR). Candidate proteins were identified by integrating differentially abundant proteins, PLS-DA features with VIP > 1 (mixOmics), and top-ranked random forest features (randomForest), with overlaps visualized using VennDiagram. Their discriminatory performance was assessed by ROC analysis and AUC using pROC. PPI networks were obtained from STRING, hub proteins were identified using PageRank, and networks were visualized in Cytoscape.

## 3. Results

### 3.1. Identification of differentially expressed tear proteins following exercise

Tear fluid samples were collected from healthy female participants both before and immediately after a standardized exercise intervention. Participant characteristics are summarized in Table S1. Following normalization, the mean protein expression levels were standardized across all samples. Principal component analysis (PCA) was conducted to evaluate intergroup variability (Figure 1A), confirming the robustness and reliability of the dataset. Proteomic data were deduplicated based on unique gene names, and proteins lacking gene annotations were excluded, yielding a total of 3173 distinct proteins identified in the tear fluid. After filtering for proteins with at least three paired pre-/post-exercise measurements, 2688 proteins were retained for subsequent differential expression analysis. In-depth analysis of protein expression changes before and after exercise revealed 484 upregulated and 260 downregulated proteins (Table S2). To visualize these findings, a volcano plot was generated for the differentially expressed proteins (Figure 1B), and a heatmap depicting the top 25 up regulated and top 25 down regulated proteins with the most significantly differential expression is presented in Figure 1C.

**Figure 1.**
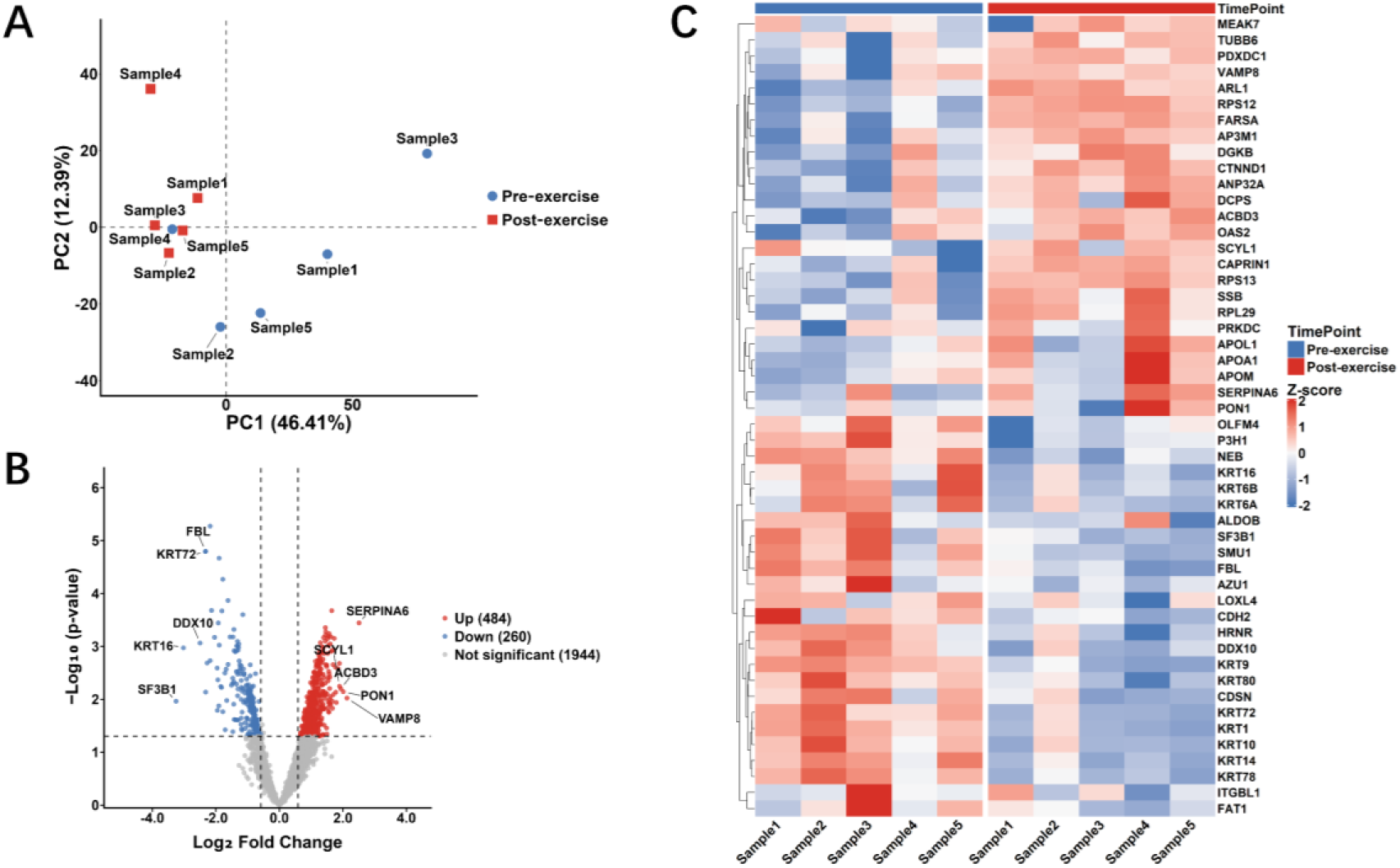
Overview of human tear protein expression in response to acute exercise. (A) PCA of tear proteomics data before and after acute exercise. (B) Volcano plot of the differentially expressed proteins. (C) Heatmap of the top 25 up-regulated and top 25 down- regulated proteins.

### 3.2. Functional enrichment and PPI network analysis of the exercise-upregulated tear proteins

To characterize the functional alterations in the tear proteome following exercise, GO and KEGG enrichment analyses were performed for the upregulated and the downregulated proteins, respectively. GO analysis showed that the exercise-upregulated proteins were predominantly associated with cytoplasmic translation, ribonucleoprotein complex biogenesis, and ribosome biogenesis. These proteins were mainly localized to cytosolic ribosomes and ribosomal subunits and primarily exhibited ribosome-related structural and RNA-binding functions (Figure 2A). KEGG analysis identified the ribosome as the most significantly enriched pathway, with endocytosis also showing notable enrichment (Figure 2B). PPI analysis identified 50 top-ranked hub proteins, most of which belonged to the ribosomal protein family (Figure 2C). Functional analysis of these hub proteins further supported their involvement in cytoplasmic translation, as structural constituents of ribosomes and components of ribonucleoprotein complexes (Figure 2D). Collectively, these results indicate that the proteins upregulated after exercise were primarily associated with protein synthesis and translational regulation.

**Figure 2.**
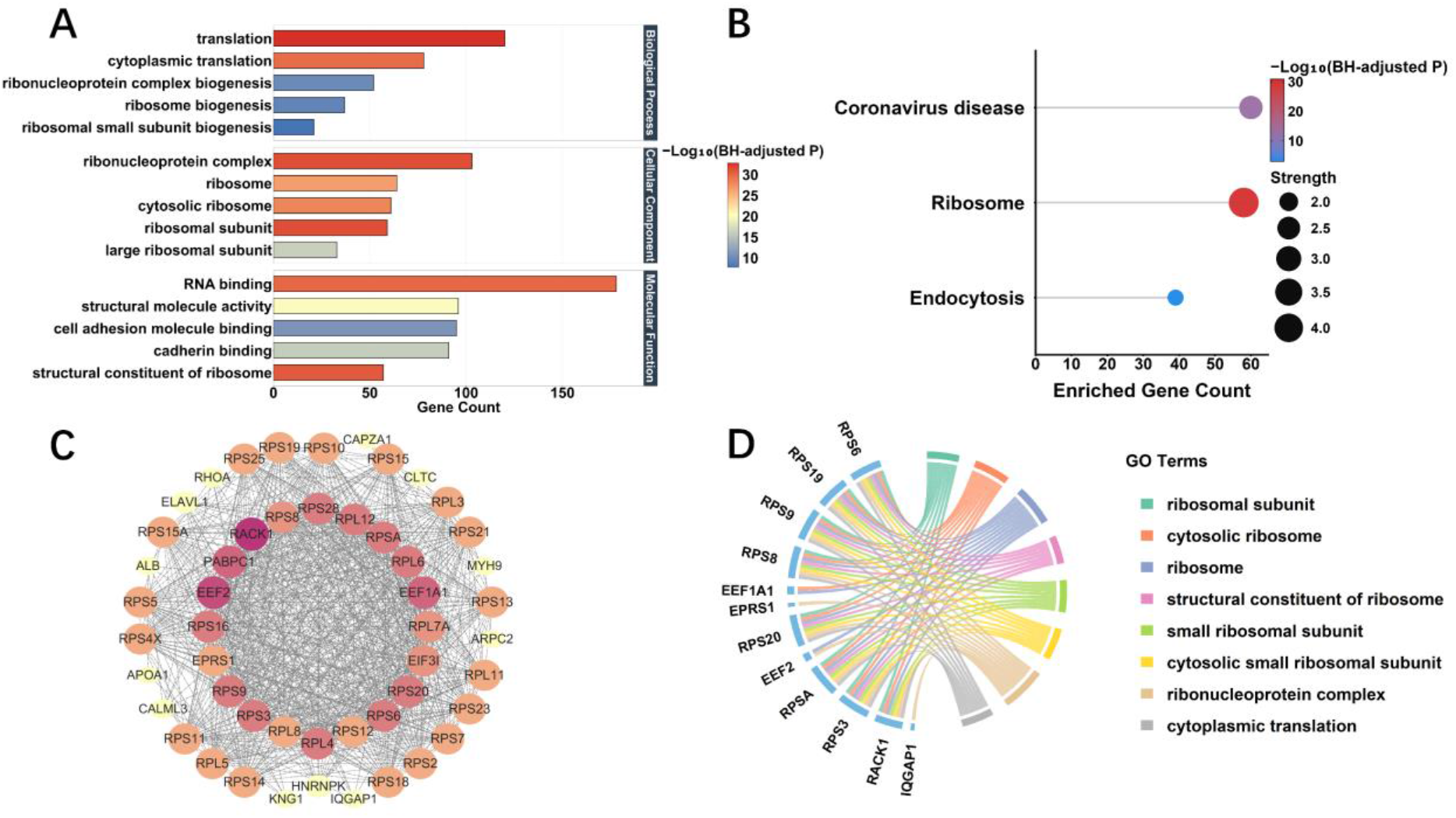
Functional enrichment and protein interaction analysis of the exercise- upregulated proteins. (A) GO enrichment bar chart of the upregulated proteins. (B) KEGG pathway enrichment analysis of the upregulated proteins. (C) PPI network of the top 50 hub proteins identified by the PageRank algorithm. (D) GO circle plot of functional enrichment results for the hub proteins.

### 3.3. Functional enrichment and PPI network analysis of exercise-downregulated tear proteins

GO enrichment analysis showed that the proteins downregulated after exercise were predominantly associated with glycoprotein metabolism, glycoprotein biosynthetic, and proteoglycan metabolism. These proteins were mainly localized to the extracellular matrix, external encapsulating structure, and lysosomal and vacuolar lumens. They also exhibited functions related to glycosyltransferase activity and extracellular matrix structural constituent (Figure 3A). KEGG analysis further identified enrichment in lysosome biogenesis, mucin-type O-glycan and glycosphingolipid biosynthesis, sphingolipid metabolism, and ECM–receptor interaction (Figure 3B). PPI analysis identified 50 top-ranked hub proteins, with EGFR, MMP9, CTSB, GLA, and GLB1 occupying central positions in the network (Figure 3C). Functional analysis of these hub proteins revealed enrichment in membrane-lipid and glycosphingolipid metabolism (Figure 3D; detailed GO terms are listed in Table S3). Collectively, these findings suggest that the proteins downregulated after exercise were primarily associated with glycosylation-related metabolism, lysosomal function, extracellular matrix organization.

**Figure 3.**
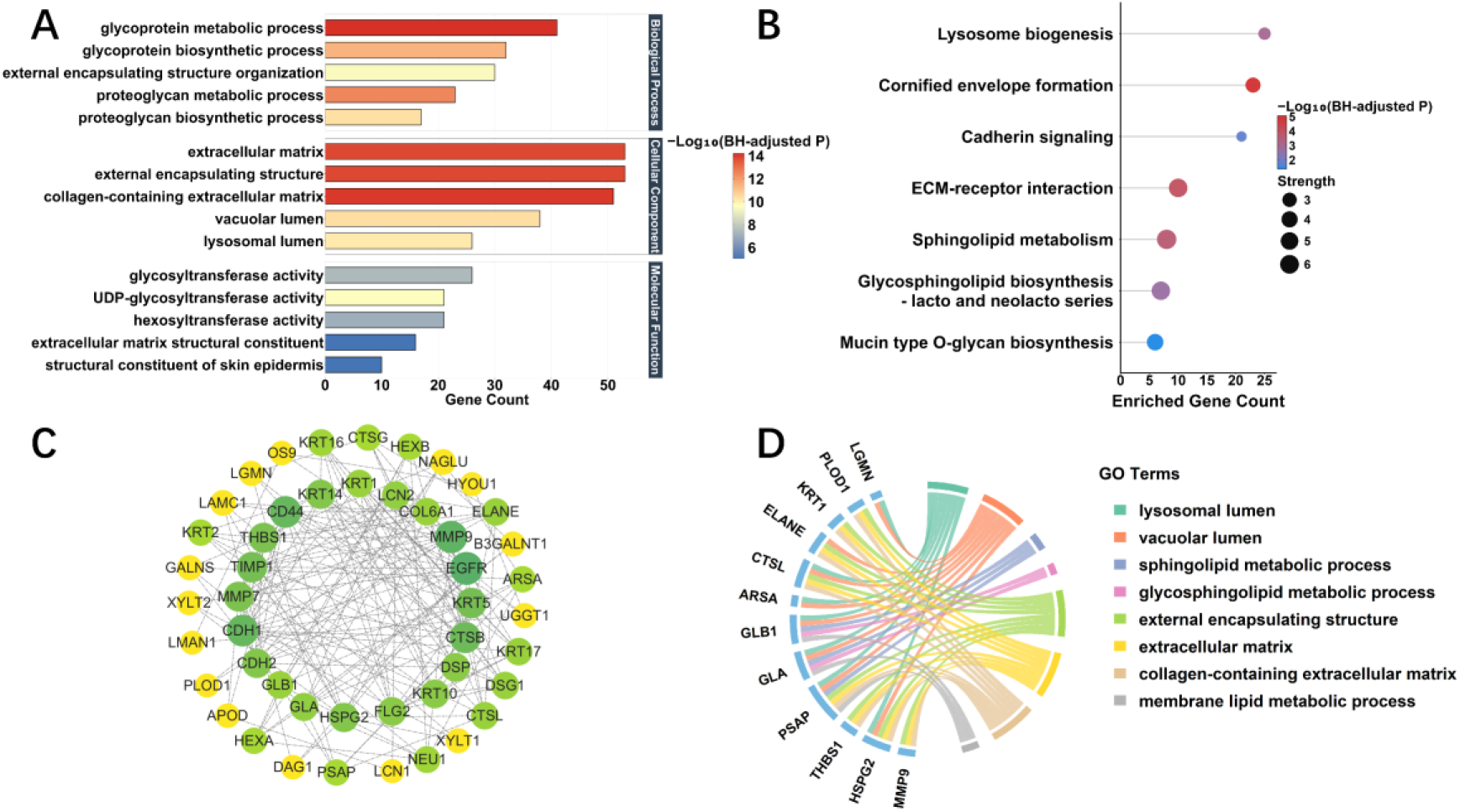
Functional enrichment and protein interaction analysis of exercise-downregulated proteins. (A) GO enrichment bar chart of the downregulated proteins. (B) KEGG pathway enrichment analysis of the downregulated proteins. (C) PPI network of the top 50 hub proteins identified by the PageRank algorithm. (D) GO circle plot of functional enrichment results for the hub proteins.

### 3.4. Gene set enrichment analysis of coordinated tear proteomic alterations following exercise

GSEA of the complete ranked protein list revealed distinct pathway-level alterations in the tear proteome following exercise. Positive enrichment was observed for MYC Targets V1 (NES = 2.492, FDR < 0.001), Mitotic Spindle (NES = 1.650, FDR = 0.033), G2m Checkpoint (NES =

1.717, FDR = 0.034), and Mtorc1 Signaling (NES = 1.584, FDR = 0.042) (Figure 4A). The enrichment of MYC target proteins was consistent with the prominent ribosomal and translational signatures identified by GO and KEGG analyses. Enrichment of the Mitotic Spindle and G2m Checkpoint gene sets indicated perturbations in cell-cycle regulation, while Mtorc1 signaling enrichment reflected changes in anabolic and translational control.

**Figure 4.**
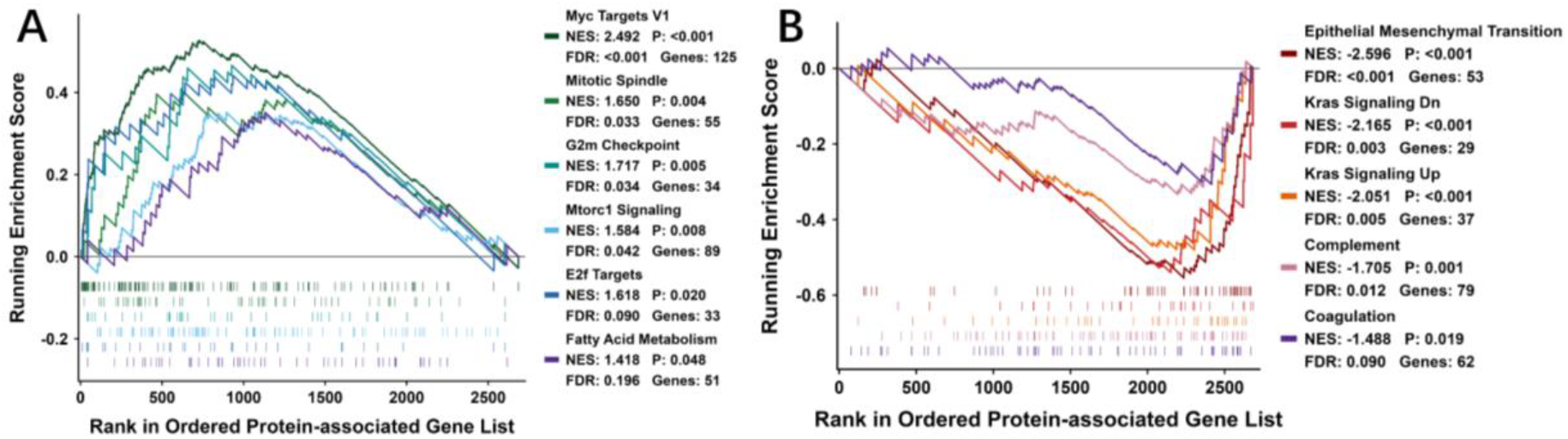
GSEA of human tear proteomic changes following exercise. (A) Positively and (B) negatively enriched Hallmark pathways after exercise. NES, normalized enrichment score; FDR, false discovery rate.

In contrast, negative enrichment was observed for Epithelial–Mesenchymal Transition (NES = −2.596, FDR < 0.001), Kras Signaling Dn (NES = −2. 156, FDR = 0.003), Kras Signaling Up (NES = −2.051, FDR = 0.005), and Complement (NES = −1.705, FDR = 0.012) were also negatively enriched (Figure 4B). These findings indicate reduced representation of proteins associated with epithelial-mesenchymal transition, Kras-related signaling, complement-mediated innate immunity, and tissue remodeling. Collectively, the GSEA results suggest that acute exercise shifted the tear proteomic profile toward enhanced translational, metabolic, and cell-cycle-adaptive signatures, accompanied by reductions in complement-immune-and tissue-remodeling-related processes.

### 3.5. Machine learning-based screening of the core tear proteins in response to exercise

Two Complementary PLS-DA and random forest analyses were applied to the 50 upregulated and the 50 downregulated hub proteins to identify candidate core proteins associated with the tear proteomic response to exercise. In the PLS-DA analysis, proteins with VIP scores greater than 1 were retained, and the 20 proteins with the highest VIP scores in each direction were selected (Figure S1A, D). Additionally, random forest models were constructed to rank protein importance (Figure 5A, C), and their stability was evaluated through 100 bootstrap resampling iterations (Figure5B, D). This analysis identified 8 upregulated and 6 downregulated proteins as stable features. Intersection of the results from the two approaches identified 7 consistently selected upregulated proteins—RPS19, RPS25, RPS23, RPS3, RPS6, RPS2 and RPS12—and 5downregulated proteins—COL6A1, GLA, GALNS, CTSL, and ARSA (Figure S1B, E).

**Figure 5.**
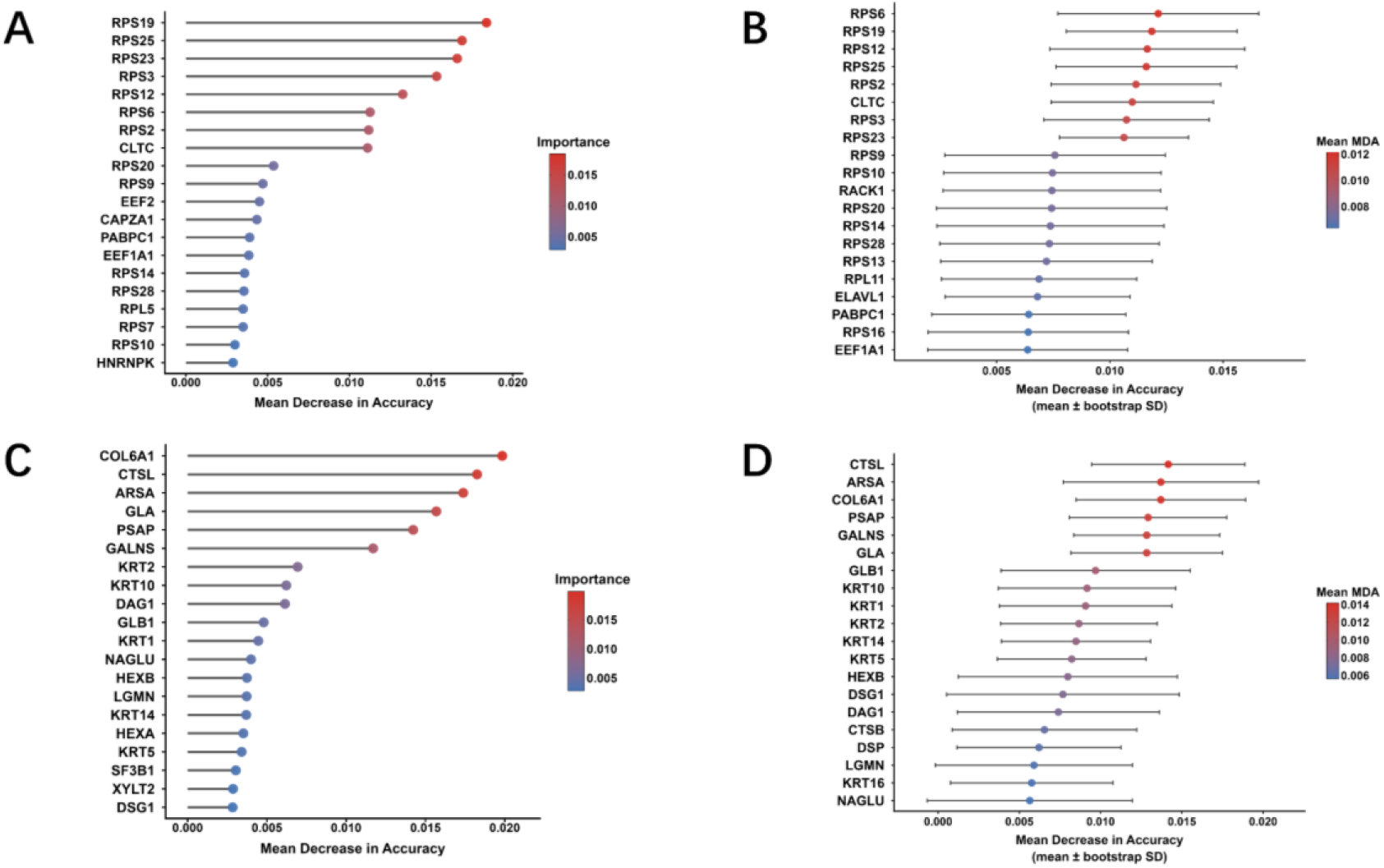
Identification of core feature proteins following exercise. (A, C) Random Forest analysis of feature proteins among upregulated and downregulated proteins, respectively. (B, D) Bootstrap stability assessment of Mean Decrease in Accuracy among upregulated and downregulated proteins, respectively.

ROC curve analysis was subsequently performed to evaluate the ability of these 12 candidate proteins to distinguish between pre- and post-exercise tear samples. Within the current dataset, each protein achieved an AUC of 1.000 (Figure S1C, F), indicating complete separation between the two exercise states. These findings identify a panel of candidate exercise-responsive tear proteins, although their discriminatory performance requires validation in larger independent cohorts.

## 4. Discussion

The present study explored the proteome of the acute exercise-induced tear, based on paired tear samples collected from healthy participants before and immediately after exercise. Among the 3,173 quantified proteins. 484 are upregulated and 260 are downregulated following exercise. These findings establish an initial molecular framework for characterizing the immediate tear response to acute exercise.

A prominent feature of the post-exercise tear proteome was the increased abundance of proteins associated with ribosomal organization and translation. GO and KEGG analyses consistently highlighted cytoplasmic translation, ribosome biogenesis, ribosomal subunits, and the ribosome pathway, with the PPI network being dominated by members of the RPS and RPL families. Positive enrichment of MYC Targets V1, Mitotic Spindle, G2m Checkpoint, and Mtorc1 Signaling was further demonstrated by GSEA. Collectively, these enriched signatures point to enhanced biosynthetic capacity and cell-cycle modulation in the tear proteome following exercise. Similar molecular responses were reported by Contrepois et al., who found that a single bout of exercise rapidly altered systemic processes related to energy metabolism, oxidative stress, growth-factor responses, and tissue repair^8^. Accordingly, the ribosomal and translational signatures detected in tears may represent a component of the rapid biosynthetic and stress-adaptive response to acute exercise.

Proteins with decreased post-exercise abundance exhibited a distinct functional profile. GO, KEGG, and PPI analyses consistently identified glycoprotein and proteoglycan metabolism, sphingolipid and glycosphingolipid metabolism, lysosomal processing, and extracellular matrix organization. Ocular surface mucins and numerous tear proteins are extensively glycosylated, and their glycan structures contribute to molecular recognition, epithelial interactions, lubrication, and ocular surface protection^24,25^. Accordingly, the reduced abundance of proteins involved in glycan synthesis and processing indicates that acute exercise rapidly modifies the biochemical composition and glycoprotein-processing profile of tears. The concurrent reduction in lysosomal enzymes, glycosphingolipid-processing proteins, and extracellular matrix-associated proteins further indicates an immediate alteration in protein degradation, membrane-component metabolism, and extracellular structural maintenance at the ocular surface.

Immune-related pathways also exhibited a coordinated downward pattern following exercise. GSEA revealed significant negative enrichment of the Complement pathway, a core component of innate immunity. In parallel, downregulation of lysosomal proteins (including lysosomal proteases such as CTSD, STSB, CTSL and LGMN) and mucin-type O-glycan biosynthesis further suggests attenuated innate immune defense and mucosal barrier function at the ocular surface. A position statement by Walsh et al. concluded that acute prolonged exercise transiently suppresses innate and mucosal immune function, creating a period of increased infection susceptibility^26^. Similarly, Gleeson et al. demonstrated that prolonged treadmill exercise reduced tear antimicrobial protein secretion rates, providing direct evidence for exercise-induced mucosal immune impairment at the ocular surface^27^. These findings align with the immune-related pathway alterations observed in the present study.

Feature-selection analyses further substantiated these functional patterns. 7upregulated proteins jointly identified by PLS-DA and random forest—RPS19, RPS25, RPS23, RPS3, RPS6, RPS2 and RPS12—are involved in ribosomal organization and translational regulation. Conversely, the 5downregulated proteins— COL6A1, GLA, GALNS, CTSL, and ARSA—participate in extracellular matrix organization, glycan modification, glycosphingolipid metabolism, and lysosomal processing. This functional correspondence with the principal enriched pathways reinforces the biological coherence of the selected protein panel. All 12 proteins achieved an AUC of 1.000 in the present dataset, indicating strong discriminatory performance within the discovery cohort; however, independent validation is required before these proteins can be considered established markers of the acute exercise response.

Previous investigations have primarily examined exercise-related changes in tear secretion, tear-film stability, oxidative stress markers, or selected inflammatory cytokines. By extending these observations to the proteome-wide level, the present study establishes an initial molecular map of the immediate tear response to exercise^28^. Tear collection is minimally invasive, requires only a small sample volume, and is amenable to repeated sampling, making it particularly suitable for assessing rapidly changing physiological states^29–31^. The functional signatures and candidate proteins identified herein provide a basis for applying tear proteomics to the investigation of individual exercise responsiveness and the development of non-invasive strategies for exercise monitoring.

Several limitations should be acknowledged. First, the small sample size and exclusive inclusion of healthy female participants limit the generalizability of the findings. Second, tear samples were collected only before and immediately after exercise, precluding characterization of the onset, peak, and recovery of the observed proteomic alterations. Third, enrichment analyses identify functional associations but do not directly quantify the corresponding biological activities within ocular surface tissues. Finally, the absence of an independent exercise-related tear proteomic dataset precluded external validation. Future investigations incorporating larger independent cohorts, multiple post-exercise time points, targeted proteomic and glycoproteomic analyses, and concurrent assessment of tear-film physiology are required to confirm these findings.

## 5. Conclusion

Acute exercise induced rapid alterations in the human tear proteome, characterized by increased ribosomal and translational signatures and decreased glycan-processing, lysosomal, and extracellular matrix signatures. This proteome-wide molecular profile establishes a preliminary reference for understanding the immediate tear response to acute exercise and provides a foundation for subsequent exercise-related tear research.

## Ethical approval

The study was approved by the Ethics Committee of the Wuhan Sports University (Approval No. 2025089)

## Data availability

All data supporting the findings of this study are included in the article and supplementary materials and can be obtained from the corresponding author upon request. Source data are provided with this article.

## Acknowledgements

This study was supported by the Natural Science Foundation of Hubei Province (2025AFD622).

## Authors’ contributions

Q.C. conceived the project and thoroughly revised the manuscript. H.Y., M.L., Q.F., J.C., and T.L., performed the human exercise experiments. M.S. analyzed the data and wrote the raw manuscript. All authors read and approved the final manuscript.

---

**Supplementary Figure 1.**
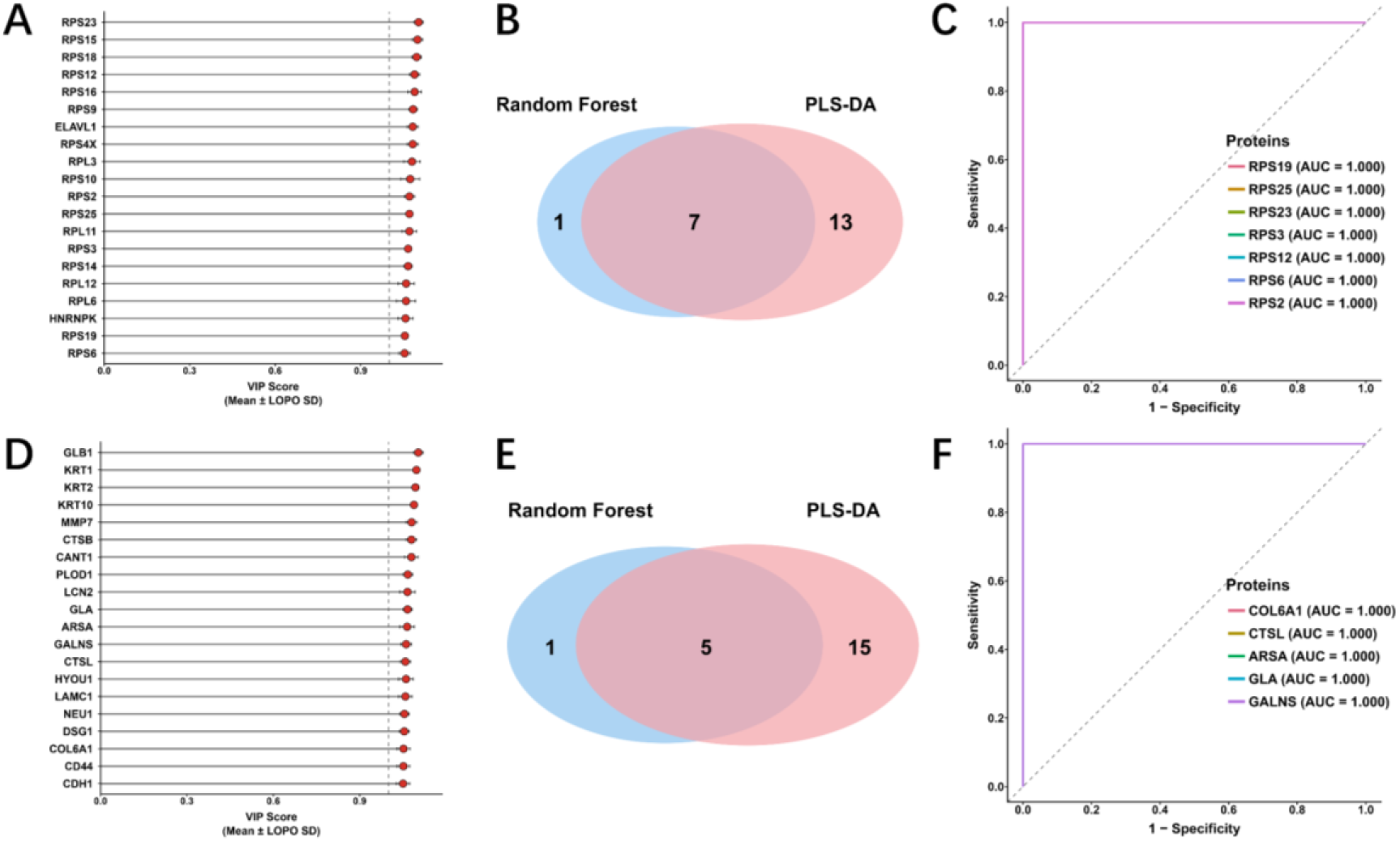
Screening and diagnostic performance of feature proteins. Upregulated proteins: (A) top 20 proteins with VIP > 1 identified by PLS-DA, (B)overlap between random forest and PLS-DA results, (E)ROC curves for overlapping proteins. **Downregulated proteins:** (C) top 20 proteins with VIP > 1 identified by PLS-DA, (D)overlap between random forest and PLS-DA results, (F)ROC curves for overlapping proteins.

